# Structural and functional analysis of VYD222: a broadly neutralizing antibody against SARS-CoV-2 variants

**DOI:** 10.1101/2025.08.28.672883

**Authors:** Meng Yuan, Brandyn R. West, William B. Foreman, Colin Powers, Ziqi Feng, Ashley L. Taylor, Xinye Yu, Grace Wang, Laura Walker, Tyler N. Starr, Robert Allen, Ian A. Wilson

## Abstract

Extensive mutations in SARS-CoV-2 spike protein have rendered most therapeutic monoclonal antibodies (mAbs) ineffective. However, here we describe VYD222 (pemivibart), a human mAb re-engineered from ADG20 (adintrevimab), which maintains potency despite substantial virus evolution. VYD222 received FDA Emergency Use Authorization for pre-exposure prophylaxis of COVID-19 in certain immunocompromised adults and adolescents. Here we show potent neutralization of this antibody against a broad range of emerging variants, including Omicron KP.3 and KP.3.1.1. X-ray crystal structures of VYD222 complexed with the receptor-binding domains of prototype SARS-CoV-2 and Omicron BA.5 demonstrate the binding epitope spans from the receptor binding site to the conserved CR3022 site. Notably, many of the matured residues between ADG20 and VYD222 occur outside the paratopic region. Deep mutational scanning indicates that SARS-CoV-2’s ability to escape VYD222 is constrained by structural compatibility and the need to maintain receptor binding. These findings provide crucial insights into the escape-resistant neutralization of VYD222 against a broad panel of clinically relevant SARS-CoV-2 variants and offer valuable guidance for risk assessment of emergent variants.

## INTRODUCTION

Since the emergence of the COVID-19 pandemic, an unprecedented effort has been directed toward the discovery and development of therapeutic antibodies against severe acute respiratory syndrome coronavirus 2 (SARS-CoV-2). However, the virus’s sustained and rapid evolution combined with antigenic shifts in the form of highly mutated emergent variants, such as Omicron BA.1 and BA.2.86, have enabled evasion from all human neutralizing antibodies (nAbs) authorized under FDA emergency use authorization (EUA). Invivyd’s discovery platform approach enabled the rapid optimization of ADG20 (also known as adintrevimab) resulting in the discovery of VYD222 (also known as pemivibart), a monoclonal antibody (mAb) with substantially enhanced breadth and potency against emergent SARS-CoV-2 variants and currently the only authorized mAb drug for SARS-CoV-2 in the U.S.

A major obstacle in the development of escape-resistant and potent therapeutic mAbs is that potency and breadth are often mutually exclusive for anti-SARS-CoV-2 nAbs—most are either potent but specific to a few variants, or broadly reactive but neutralize many variants with modest to low potency (*1*). For example, mAbs targeting the variable receptor-binding site (RBS) directly impede the spike protein from engaging with the host receptor. These RBS antibodies tend to neutralize with higher potency, but often at the expense of reduced neutralizing breadth due to high variation in the RBS (Figure 1A-B). Conserved epitopes on SARS-CoV-2 spike generally do not overlap with the RBS and generally tend to elicit antibodies with lower potency, which likely explains why neutralizing potency and breadth are often somewhat mutually exclusive. Interestingly, we and others identified a novel epitope that spans from one end of the RBS to the highly conserved CR3022 site (*2*). Several broad-and-potent nAbs have been discovered targeting this RBS-D/CR3022 or class 1/4 site including SA55 and SC27 (*1, 3–5*). We previously isolated a broad mAb (ADI-55688) targeting this site from a SARS-CoV-1-convalescent donor (*6*) that was subsequently affinity matured against SARS-CoV-2 resulting in ADG20. ADG20 demonstrated a nearly 200-fold improved binding affinity compared to ADI-55688 and exhibited extraordinary potency and neutralization breadth across the sarbecovirus genus. However, ADG20 demonstrated reduced activity to the antigenically distinct Omicron BA.1 variant and potency was further reduced against BA.2 and descendent variants (*1, 7, 8*).

**Figure 1.**
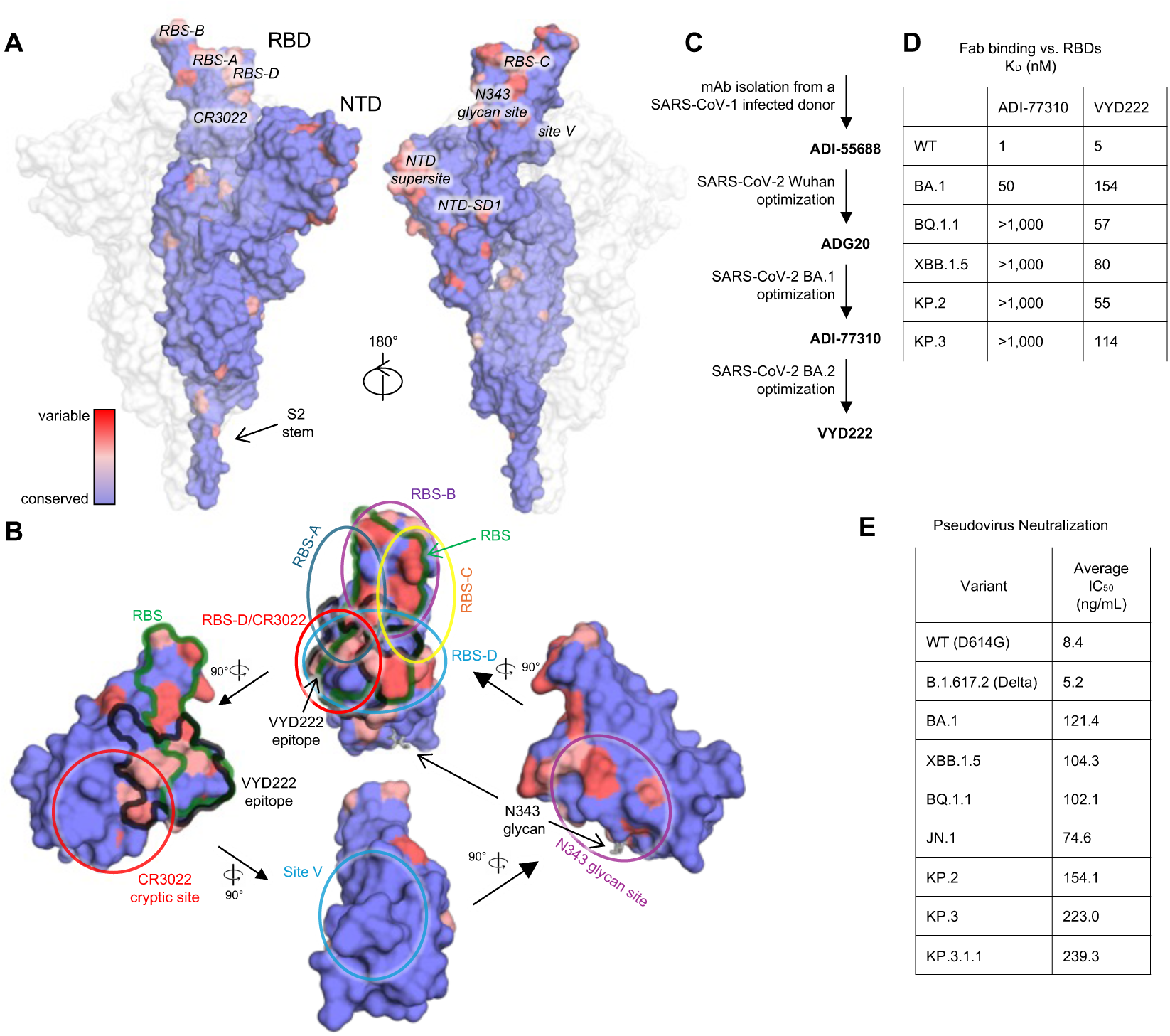
Overview of vulnerable sites to antibodies on SARS-CoV-2 spike. (A-B) Sequence conservation of **(A)** spike proteins of SARS-CoV-2 variants and **(B)** RBD was calculated by ConSurf (*38*). Epitopes are labeled at their corresponding positions. The color scale between panels A and B is consistent. **(B)** The VYD222 epitope and RBS are highlighted with black and green outlines, respectively. The N343 glycan is shown as sticks. Antibody target sites on the RBD are categorized into seven sites, namely, RBS-A, -B, -C, -D, CR3022 cryptic site, N343 glycan site (*1*), and site V (*43*), are indicated by circles. **(C)** Engineering of the ADI-55688-lineage antibodies that led to VYD222. **(D)** Binding affinities of Fabs and SARS-CoV-2 WT and variant RBDs measured by biolayer interferometry (BLI). **(E)** Neutralization of VYD222 against pseudotyped SARS-CoV-2 variants. Average IC_50_ values for VYD222 pseudovirus neutralization in the PhenoSense assay are listed.

Here, we report on a therapeutic mAb re-engineered from ADG20 to enhance and broaden neutralization activity against Omicron and Omicron sub-lineages in two steps. Step 1 involved affinity maturation of ADG20 against Omicron BA.1 which yielded ADI-77310 with high potency against Omicron BA.1. Step 2 involved the affinity maturation of ADI-77310 against BA.2 resulting in the isolation of ADI-80222 that demonstrated exceptional neutralization breadth and potency against a wide range of clinically relevant SARS-CoV-2 sub-lineages. ADI-80222 was further engineered in the Fc domain to include the LA half-life extending modification resulting in VYD222, which is otherwise indistinguishable in the Fab domain compared to ADI-80222 rendering the two mAbs identical in the context of the structural data presented here. VYD222 was granted EUA in the United States in March 2024 for prevention of symptomatic COVID-19 in certain adults and adolescents (aged ≥12 years and weighing >40 kg) with moderate-to-severe immunocompromise who are unlikely to mount adequate immune response to COVID-19 vaccination (*9, 10*). In this study, we present an in-depth structural analysis using x-ray crystallography, revealing atomic interactions between ADI-77310 and VYD222 with the receptor-binding domains (RBDs) of the prototype SARS-CoV-2 and the BA.5 variant. These structures offer a detailed explanation of the expanded neutralization breadth of VYD222 and its ability to accommodate mutations arising from viral evolution. Additionally, we employed deep mutational scanning (DMS) to identify potential RBD mutations that could reduce VYD222 binding. Overall, our comprehensive structural and functional analysis of VYD222 uncovers the mechanisms behind its extraordinary neutralization activity and breadth, offering clinically relevant critical insights for the assessment of VYD222 activity against emergent SARS-CoV-2 variants.

## RESULTS

### VYD222 potently neutralizes early and circulating SARS-CoV-2 variants

The therapeutic mAb VYD222 is an Fc-modified version of ADI-80222, which was derived from multiple rounds of yeast-display affinity maturation of a human anti-SARS-CoV-1 mAb ADI-55688 with intermediate mAbs ADG20 (*1*) and ADI-77310 (Figure 1C). VYD222 contains M428L and N434A mutations on the constant region of the immunoglobulin G1 (IgG1) to extend its half-life. VYD222 strongly binds to all tested SARS-CoV-2 variants (Figures 1D and S1). The neutralization breadth and potency of VYD222 were evaluated against a broad range of SARS-CoV-2 pseudoviruses, including early variants D614G, Delta, BA.1 and XBB.1.5 with mean IC_50_ values ranging from 5.2 to 121.4 ng/ml. Although VYD222 was only affinity matured against BA.1/BA.2, it retained potent neutralizing activity against currently circulating variants including KP.3 and KP.3.1.1 (neutralization IC_50_ values 223.0 and 239.3 ng/ml, respectively) (Figures 1E and S2).

### Crystal structures of ADI-77310 and VYD222 in complex with SARS-CoV-2 RBD

We determined three crystal structures of the re-engineered antibodies: ADI-77310 in complex with the prototype RBD (RBD-WT), and VYD222 in complex RBD-WT and BA.5 RBD, at resolutions of 3.06 Å, 3.00 Å, and 2.70 Å, respectively (Table S1 and Figure 2A). As for the ancestral antibodies ADI-55688 and ADG20 (*1*), ADI-77310 and VYD222 also target the RBD at the corner distant from the ridge region with the same angle of approach and through interaction with CDRs H1, H2, H3, L1, and L3 (Figure S3A). VYD222 targets BA.5 RBD in the same binding pose and epitope as the RBD-WT (Figures 2A and S3A). We calculated the buried surface area (BSA) of 210 RBD-targeting antibodies (Table S2) from their structures in the Protein Data Bank (PDB) using PISA (*11*). The average BSA was 904 Å^2^ with a standard deviation of 151 Å^2^. The BSAs on the RBD-WT buried by ADI-77310 and VYD222 are 718 Å^2^ and 701 Å^2^, respectively, and that on the BA.5 RBD buried by VYD222 is 745 Å^2^, demonstrating that the ADG20/VYD222 lineage of antibodies has a smaller binding interface on average than RBD-targeting antibodies. The epitopes of ADI-77310 and VYD222 overlap with the receptor binding site (RBS) and therefore likely have a mechanism of action resulting from directly impeding binding of the RBD to the host receptor ACE2 (Figure 2). However, their epitopes are only exposed when the RBDs are in the up conformation (Figure S3B).

**Figure 2.**
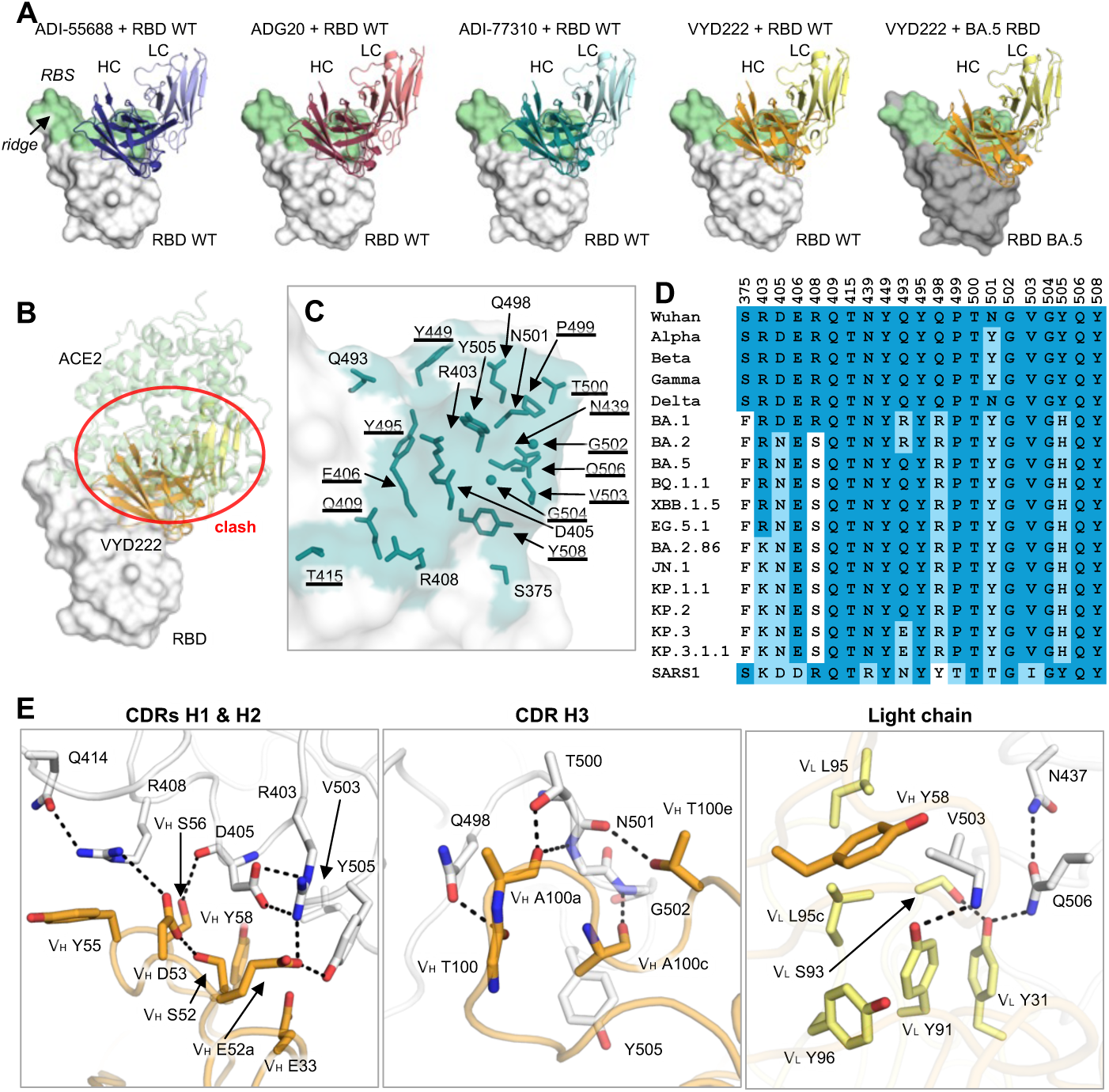
Crystal structures of VYD222-lineage antibodies in complex with SARS-CoV-2 RBD. **(A)** Overall view of crystal structures of RBD in complex with the lineage of antibodies derived from ADI-55688. WT and BA.5 RBDs are represented by surfaces in white and dark grey, respectively. Antibodies ADI-55688, ADG20, ADI-77310, and VYD222 are shown in blue, salmon, teal, and orange/yellow colors with heavy and light chains in dark and light colors. For clarity, only the variable domains of the antibodies are shown. Structures of ADI-55688 and ADG20 were determined and reported in our previous study (*1*) (PDBs 7U2E and 7U2D) and are shown here for comparison. The receptor binding site (RBS) is highlighted in light green. Epitope residues and RBS residues are defined by residues with surface area buried by the antibodies and ACE2 of >0 Å^2^, respectively, and calculated based on the antibody/RBD crystal structures and an RBD/ACE2 complex structure (PDB 6M0J) (*44*) by PISA (*11*). The definition of epitope and RBS residues, as well as the color coding for the antibodies and RBDs are consistent in this paper. **(B)** The ACE2/RBD structure superimposed on the VYD222/RBD structure. VYD222 would clash with ACE2 when bound to RBD (indicated by red circle). **(C)** Epitope residues of VYD222 on RBD WT. Residues that are conserved across all the major SARS-CoV-2 variants as shown in panel D are underlined. **(D)** Sequence alignment of epitope residues of VYD222. Identical residues are highlighted with a darker blue background, while similar residues [amino acids scoring ≥ 0 in the BLOSUM62 alignment score matrix (*45*)] to SARS-CoV-2 WT have a light blue background, and dissimilar residues are shown with a white background. SARS-CoV-1 is shown here for comparison as the ancestral antibody, ADI-55688, was isolated from a SARS-CoV-1-infected donor. **(E)** Detailed molecular interactions between VYD222 CDRs and RBD WT. Hydrogen bonds and salt bridges are represented by dashed lines.

The paratope residues of the heavy chain of VYD222 form extensive interactions with the WT RBD (Figure 2E). V_H_ S52 hydrogen bonds (HBs) with V_H_ D53 where the latter forms a salt bridge with RBD-R408. R408 is further stabilized by a cation–π interaction with V_H_ Y55. V_H_ S56 HBs with the mainchain carbonyl group of RBD-D405, and the side chain of D405 forms two salt bridges with R403. V_H_ E52a also participates in the network of interactions by forming a salt bridge RBD-R403 and an HB with Y505. Hydrophobic interactions between V_H_ Y58 and RBD-V503 also stabilize the interaction. The mainchain carbonyl groups of V_H_ T100, A100a, and A100c HB with RBD-Q498, T500, N501, and G502. The sidechain hydroxyl group of V_H_ T100e HBs with RBD-N501. The side chain of V_H_ A100c makes hydrophobic interactions with RBD-Y505 carbonyl oxygen. The light chain of the antibody also participates in the interaction with the antigen. V_L_ S93, V_L_ Y31, RBD-Q506, and N437 form a series of tandem HBs. The sidechain of the highly conserved RBD-V503 occupies a hydrophobic pocket comprising of V_L_ Y91, L95, L95c, Y96, and V_H_ Y58. V_L_ Y91 also HBs with the mainchain amide of V503. Although several residues of the antibody have mutated during cycles of antibody re-engineering, 21 out of the 24 paratope residues of VYD222 remain unchanged from the ancestral antibody ADI-55688 (Figure 3A). These residues compose the RBD recognition nucleus of this lineage of antibodies.

**Figure 3.**
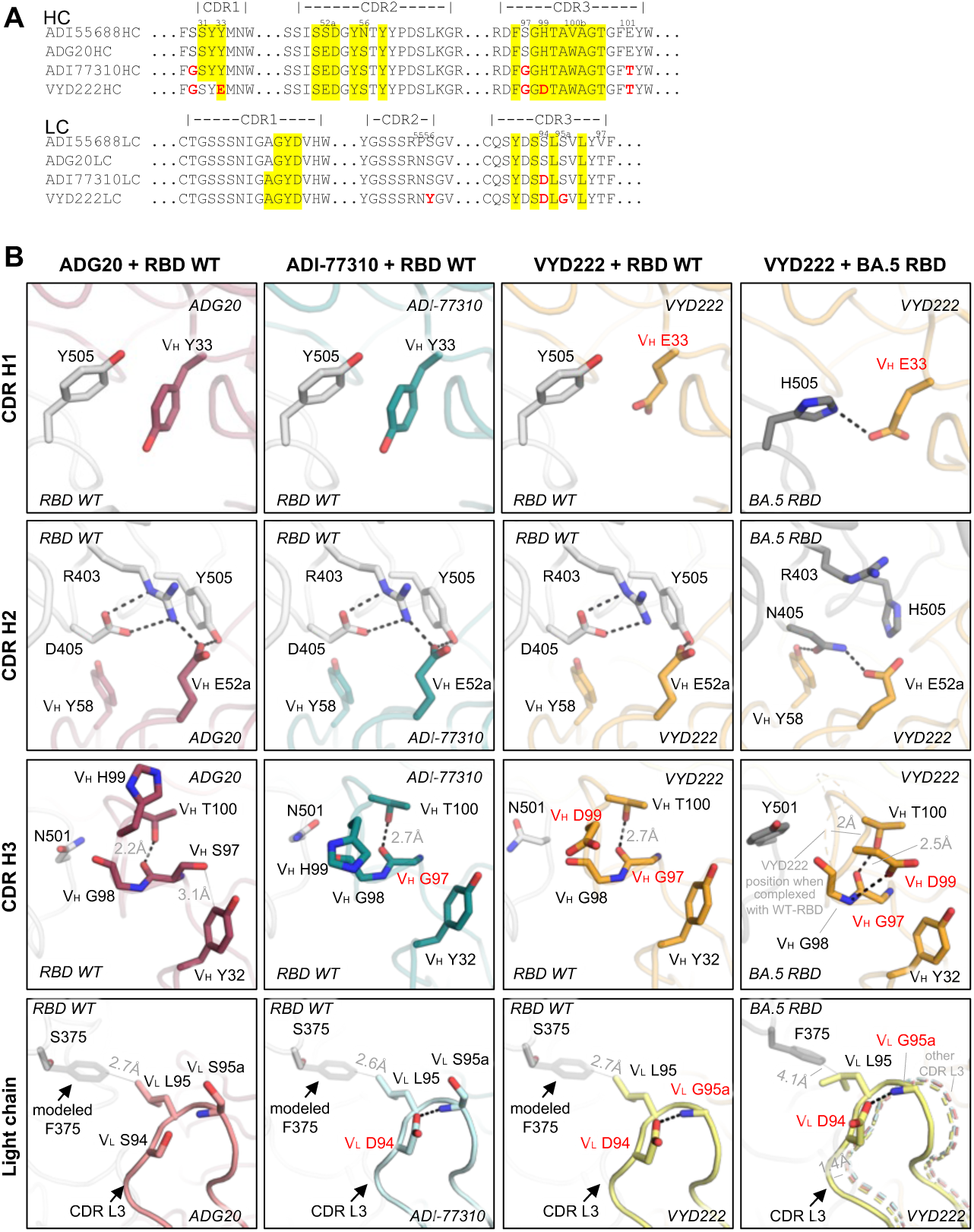
Structural basis of the broadened antibody reactivity. **(A)** Sequence alignment of the lineage of antibodies derived from ADI-55688. Residues of ADI-77310 and VYD222 that differ from ADI-55688 and ADG20 are highlighted in red. Paratope residues of VYD222 (surface area buried by antigen > 0 Å^2^ as calculated by PISA (*11*) using the antibody/RBD-WT structures) are highlighted in a yellow background. Location of the CDR loops (Kabat numbering) are indicated on top of the sequences. **(B)** Detailed interactions between the antibodies (ADG20, ADI-77310 and VYD222) and antigens (WT and BA.5 RBDs). Hydrogen bonds and salt bridges are represented by black dashed lines. In some panels, loops from another structure are represented by thin, dashed cartoon for structural comparison.

### Structural basis of restored neutralization against emerging variants

The binding and neutralization activity of ADG20 was greatly reduced against Omicron BA.1 (*1*) but restored by mutations in antibody ADI-77310 (Figures S1, 1D and 3A). Additional mutations in VYD222 facilitated potent neutralization to all tested SARS-CoV-2 variants, including BA.1, BA.5, XBB.1.5, KP.2, and KP.3.1.1 (Figures 1D-E and S2) (*12*). Notably, most of the mutated residues are not paratopic residues—none of the four mutations of ADI-77310 (on top of ADG20) is a paratopic residue, and only two out of eight mutations in VYD222 (on top of ADG20) locate to the paratope (Figure 3A). In other words, non-paratope mutations play a vital role in enhancing the antibodies’ potency and breadth, as observed in some previously reported mAbs (*13–15*). To investigate the structural basis of affinity restoration against emerged variants, we systematically compared the structures of ADG20, ADI-77310, VYD222 complexed with RBD-WT, and VYD222 with BA.5 RBD (Figure 3B).

The CDR H1 of ADI-77310 remains unchanged compared to ADG20, where V_H_ Y33 stacks with RBD-Y505 (Figure 3B). Since Omicron BA.1, Y505 was mutated to H505 (Figure 2D). The paratope mutation V_H_ Y33E in VYD222, in comparison to ADI-77310, now forms a salt bridge/HB to the H505 RBD mutation in BA.5 (Figure 3B). Residues on CDR H2 are conserved from ADG20 through VYD222. In all three structures of ADG20, ADI-77310, and VYD222 complexed with RBD-WT, the RBD-D405 form two salt bridges with R403 that help align it with Y505 to form a network of interactions with the antibodies’ paratope residue V_H_ E52a. Although two out of the three epitope residues on the RBD were mutated from the BA.2 subvariant (i.e. D405N and Y505H), these residues change their rotamers and form a different set of interactions with VYD222 as shown in the BA.5 panel (Figure 3B). In addition to the interaction between RBD-H505 and V_H_ E33, RBD-N405 makes two HBs with V_H_ E52a and Y58, respectively. In CDR H3, ADG20 exhibits a slightly unfavorable internal clash where V_H_ S97 is crowded by V_H_ Y32 so that its mainchain carbonyl is only 2.2 Å from V_H_ T100. This conflict is resolved by the non-paratopic V_H_ S97G mutation in ADI-77310 and VYD222. The lack of a sidechain in V_H_ G97 allows more room, and then a more favorable HB (2.5–2.7 Å) between V_H_ G97 and T100. The N501Y mutation emerged in late 2020 and has dominated SARS-CoV-2 variants from late 2021 to the present (Figure S3C). CDR H3 of VYD222 accommodates N501Y, where the backbone of the CDRH3 loop shifts by 2 Å upon binding to the mutated antigen and accommodates the bulkier amino acid.

The CDR L3 also extensively interacts with the RBD, especially with the heavily mutated region 371-375 arising from Omicron BA.1 (S371L/F, S373P, and S375F). The mutation V_L_ S95aG, which separates VYD222 from ADI-77310, enhances the flexibility of the CDR L3 loop, which allows CDR L3 to accommodate S375F. In fact, the crystal structure of VYD222 with F375-containing BA.5 RBD demonstrates a 1.4-Å shift of CDR L3, resulting in a favorable hydrophobic interaction between V_L_ L95 and RBD-F375 (distance 4.1 Å).

Overall, the structural analysis explains the restoration of neutralization activity against emerging variants by the mutated residues in VYD222, where V_H_ Y33E restores interaction with RBD-Y505H, V_H_ S97G and H99D provide more accommodation for RBD-N501Y, and mutations in the light chain enhance the flexibility of CDR L3 for interaction with S375F. In short, not only paratope but also non-paratope mutations play important roles in enhancing antibody binding to SARS-CoV-2 variants.

### Deep mutational scanning of viral escape from VYD222

To evaluate potential viral escape of VYD222, we performed a deep mutational scan for VYD222 against various SARS-CoV-2 variants (Figures 4 and S4). We determined the impact of all single amino acid mutations in the context of Wuhan-Hu-1, Omicron BA.2, XBB.1.5, and BA.2.86 RBDs on VYD222 binding (Figure 4A). Mutations of epitope residues 403-405, 407-409, 436-437, and 500-506 exhibit escape from the antibody. However, most of these mutations result in substantial reduction of receptor binding, which may therefore decrease viral fitness. In contrast, potentially compatible mutations, as suggested by a sequence conservation analysis of ACE2-binding sarbecoviruses and SARS-CoV-2 variants (Figure 4B), usually exhibit limited escape against VYD222. For example, mutation R403K in subvariants of BA.2.86 is well accommodated by VYD222 (Figure 4A). Residue 403 is not exclusively Arg or Lys among all ACE2-binding sarbecoviruses. An ACE2-binding sarbecovirus, RaTG13, contains T403, suggesting that a Thr might be compatible with SARS-CoV-2 (Figure S5). The crystal structure of VYD222/BA.5 RBD shows that a basic R403 (or K403) is accommodated in a negatively charged region of VYD222, while mutation to T403 would reduce this electrostatic interaction (Figure 4C). This is in line with the R403T mutation retaining ACE2 binding while conferring a modest degree of escape from VYD222 binding, as observed in our DMS experiment (Figures 4A and 4D). Likewise, at position 504, an Asn is found in the Pangolin_GXP2V strain despite the dominance by G504 in almost all sarbecoviruses (Figures 4B and S5). G504N exhibits minimal binding loss in the DMS experiment (Figure 4D). A VYD222/BA.5 RBD-G504N structural model predicted by FoldX suggests that N504 can be accommodated by VYD222 (Figure 4E) with a slightly reduced antibody-antigen binding energy (ΔΔG_binding_ = 1.5 kcal/mol). RBD residues in the T500-Q506 loop extensively interact with VYD222 (Figure 4F), and mutations of this loop result in escape from VYD222 (Figure 4A). Notably, escape variants carrying spike substitutions at T500 were observed both following serial passage of Omicron variants in the presence of VYD222 in cell culture and in participants treated with VYD222 in the CANOPY clinical study (*9*). However, T500, as well as G502, and Q506 in the T500-Q506 loop, are exclusively conserved among all ACE2-binding sarbecoviruses, which may suggest other potential functional constraints of these residues in nature (Figure 4B). Mutations at positions 501, 503, and 505 are either well accommodated by VYD222 or result in a significant loss of receptor binding, suggesting the limited escapability of these residues. Overall, our findings demonstrate that SARS-CoV-2’s ability to escape from VYD222 is constrained by its structural compatibility and the necessity of maintaining receptor binding.

**Figure 4.**
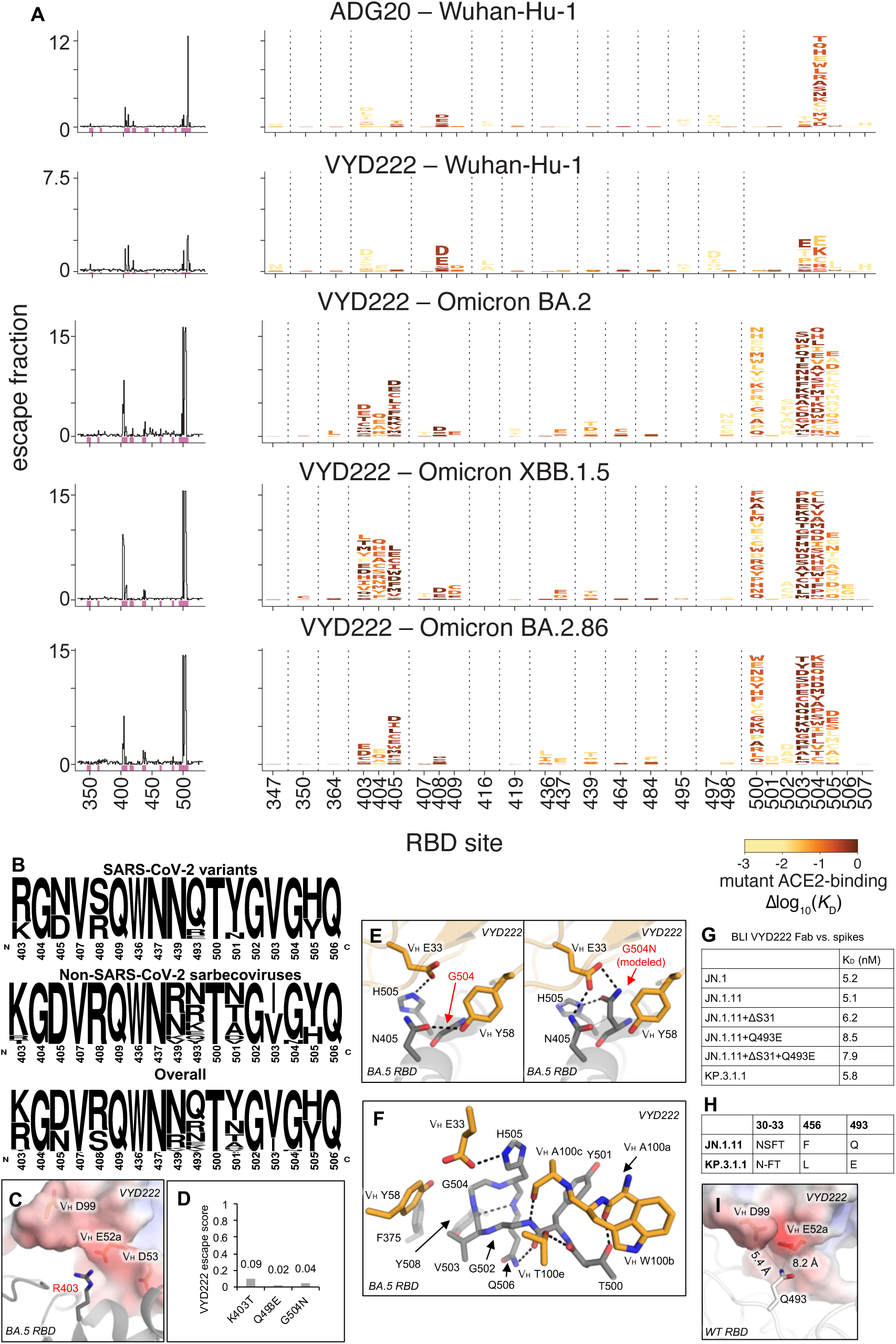
Limited escapability of SARS-CoV-2 from VYD222. **(A)** Deep mutational scanning maps of mutations that escape binding by ADG20 and VYD222. Sites with strong escape mutations are shown in the logo plots, where taller amino acid letters represent stronger escape-causing mutations. Amino acid letters are colored according to how deleterious mutations are for ACE2 binding. See Figure S4 for experimental details. **(B)** Sequence conservation of the VYD222 epitope among ACE2-binding sarbecoviruses. Amino acid residues from ACE2-binding sarbecoviruses as identified in Starr et al. 2022 (*46*) and SARS-CoV-2 variants were used for this analysis by WebLogo (*47*). Strains used for this analysis are listed in Figure S5. **(C)** Interactions between VYD222 and BA.5 RBD-R403. **(D)** Escape scores of K403T, Q493E, and G504N against VYD222 in the context of BA.2.86. An escape score of 1 represents complete escape while 0 is no escape. R403 was mutated to K in BA.2.86 and later variants. **(E)** Interactions between VYD222 (orange) and RBD-G504. Structures of the mutated RBDs are modeled by FoldX (*48*) and shown in the right panel. **(F)** Interactions between VYD222 (orange) and the 500-505 loop of RBD. **(G)** Binding affinities of VYD222 Fab and SARS-CoV-2 spike proteins measured by BLI. **(H)** Sequence differences between SARS-CoV-2 variants JN.1.11 and KP.3.1.1. **(I)** Interactions between VYD222 and RBD-Q493. The electrostatic surface potential of VYD222 is calculated by APBS (*49*). Hydrogen bonds and salt bridges are represented by black dashed lines.

The SARS-CoV-2 variant KP.3 carries a Q493E mutation, while KP.3.1.1 has an additional S31 deletion in ^30^NSFT^33^, creating a N-linked glycosylation site at the ^30^NFT^33^ motif as demonstrated in our previous mass spectrometry analysis (*16*). Our single-mutagenesis study showed that the Q493E mutation led to a modest reduction in VYD222 binding compared to the JN.1.11 variant as well as JN.1.11+ΔS31 (Figures 4G-H and S6), which correlates with the low escape score for Q493E observed in our DMS analysis (Figure 4D). The sidechain of RBD-Q493 is typically accommodated by a negatively charged region on VYD222, which may be less favorable for the acidic Q493E mutation, explaining the very slightly reduced binding affinity (Figures 4I and S6). We also assessed VYD222 binding against spike proteins carrying the S31 deletion, as well as the double mutant (ΔS31+Q493E), both of which showed slightly reduced binding (Figures 4G and S6). KP.3.1.1 also contains both mutations and binds strongly to VYD222, consistent with the potent neutralization observed for VYD222 against KP.3.1.1 (Figure 1E).

## DISCUSSION

Receptor binding sites (RBSs) are relatively conserved regions in many viruses, such as those present in the HIV envelope protein (Env) (*17*) and, to a lesser extent, influenza hemagglutinin (HA) (*18, 19*) and, because these regions are critical for binding to host cell receptors, mutations in the RBS could disrupt this essential function. In contrast, the RBS of SARS-CoV-2 has demonstrated substantial plasticity with many variant lineages incorporating mutations in the RBS often resulting in strong escape from nAbs elicited by natural infection or vaccination, while maintaining or even increasing ACE2 receptor binding.

Despite the plasticity demonstrated by the SARS-CoV-2 RBS, this region remains an attractive target for antibodies, which often achieve potent neutralization via direct inhibition of receptor binding. Thus, it is not surprising that a majority of SARS-CoV-2 clinical antibodies target one of the four epitopes previously characterized by our group: RBS-A, RBS-B, RBS-C, and RBS-D (*1, 20*) (Figure 1A-B). The RBS-A and RBS-B epitopes [related to class 1 and class 2 in (*21*)] are targeted by public antibody classes encoded by IGHV3-53/3-66, such as Etesevimab/CB6 and Amubarvimab/P2C-1F11 (RBS-A) (*22, 23*) or Casirivimab/REGN10933 and Regdanvimab/CT-P59 (RBS-B) (*24, 25*). Public antibodies encoded by IGHV1-2 (*20, 26*) and IGHV1-58/IGKV3-20 (e.g. Tixagevimab/COV2-2196) (*27*) also target the RBS-B epitope. The RBS-C epitope is targeted by IGHV1-69 antibodies (Bamlanivimab/LY-CoV555) (*28*), while RBS-D is targeted by Imdevimab/REGN10987 (*24*), Cilgavimab/COV2-2130 (*27*), and Bebtelovimab/LY-CoV1404 that belongs to the IGHV2-5/IGLV2-14 encoded public class (*29, 30*). The four epitopes are highly variable and exhibited viral escape from all of the clinically relevant nAbs described above that target these sites.

However, we previously identified a more general site accommodating broad and potent nAbs, namely the RBS-D/CR3022 site (*1*) [corresponding to ‘epitope F’ by Cao et al. (*31*)]. This site spans from one end of the RBS to the top end of the highly conserved CR3022 site. These mAbs were found to be highly potent against a wide range of SARS-CoV-2 variants as well as other sarbecoviruses. The SARS-CoV-2 virus successfully introduced multiple mutations at this site simultaneously including S371L, S375F, Q493R, Q498R, N501Y, and Y505H, in the context of the Omicron B.1.1.529 variant which emerged in December of 2021. These mutations were further combined with an additional set of mutations introduced by the BA.2 variant (D405N, and R408S, as well as S371F instead of S371L in BA.1) which emerged in January of 2022, and together these mutations contributed to escape the majority of antibodies that bound to this region, which has also carried through to contemporary emergent variants.

Here, we introduce and characterize a broad and potent nAb, VYD222, which binds to the RBS-D/CR3022 site, similar to its ancestral mAb ADG20 and the related SARS-CoV-1 mAb ADI-55688. VYD222 was successfully engineered using the Invivyd discovery platform to expand upon the breadth and potency of ADG20 to further include Omicron and descendant lineages while maintaining the favorable characteristics of the RBS-D/CR3022 epitope. Notably, our DMS studies demonstrate that many of the VYD222 binding escape mutations that occur on the structure-defined VYD222 epitope result in substantial reduction of receptor binding and are, therefore, unlikely to emerge in the context of a fit variant. In contrast, RBD mutations within the VYD222 epitope, which are most compatible with receptor binding, typically exhibited limited escape against VYD222. Furthermore, despite multiple rounds of affinity maturation, the paratope remains highly conserved with only 4 differences between VYD222 and the original human-derived SARS-CoV-1 mAb ADI-55688 out of 24 total paratope residues involved in the binding interface, highlighting framework residue modifications in enhancing mAb potency and breadth.

The structural characterization of the VYD222 epitope combined with neutralization and escape data from DMS studies enables a thorough risk assessment of emergent variants. While there remains a small possibility of escape from mutations acting distal to the epitope that are currently unknown or unappreciated, the most likely scenario for viral escape is the incorporation of a non-tolerable mutation directly in the VYD222 epitope. Our combined structural and functional data therefore enable a risk assessment of emergent variants with respect to the VYD222 epitope. For example, an emergent variant containing one or more significant DMS escape mutations located within the structurally defined epitope are more likely to threaten the activity of VYD222. Likewise, emergent variants that do not contain mutations proximal to the epitope are unlikely to significantly impact VYD222 activity.

Overall, our work demonstrates the great potential of the RBS-D/CR3022 site for developing broad and potent antibodies, such as VYD222, which are resistant to immune escape, while providing a framework for the risk assessment of emergent variants with respect to VYD222 activity.

## MATERIALS AND METHODS

### Expression and purification of recombinant proteins

The heavy and light chains of antibody Fabs and IgGs were cloned into phCMV3. The plasmids were transiently co-transfected into ExpiCHO cells at a ratio of 2:1 (HC:LC) using ExpiFectamine™ CHO Reagent (Thermo Fisher Scientific) according to the manufacturer’s instructions. The supernatant was collected at 10 days post-transfection. The IgGs and Fabs were purified with a CaptureSelect™ CH1-XL Affinity Matrix (Thermo Fisher Scientific) followed by size exclusion chromatography.

Expression and purification of the SARS-CoV-2 spike receptor-binding domain (RBD) for crystallization were as described previously (*2*). Briefly, the RBDs (residues 333-528) of the SARS-CoV-2 spike (S) protein (GenBank: QHD43416.1) and the Omicron BA.1, BA.5, BQ.1.1, XBB.1.5, XBB.1.16, JN.1, KP.2, and KP.3 S were cloned into a customized pFastBac vector (*32*), and fused with an N-terminal gp67 signal peptide and C-terminal His_6_ tag (*2*). A recombinant bacmid DNA was generated using the Bac-to-Bac system (Life Technologies). Baculovirus was generated by transfecting purified bacmid DNA into Sf9 cells using FuGENE HD (Promega), and subsequently used to infect suspension cultures of High Five cells (Life Technologies) at an MOI of 5 to 10. Infected Sf9 cells were incubated at 28°C with shaking at 110 r.p.m. for 72 h for protein expression. The supernatant was then concentrated using a 10 kDa MW cutoff Centramate cassette (Pall Corporation). The RBD protein was purified by Ni-NTA, followed by size exclusion chromatography, and buffer exchanged into 20 mM Tris-HCl pH 7.4 and 150 mM NaCl.

All spike ectodomain constructs contain a C-terminal T4 fibritin trimerization domain, and a Twin-strep-tag for purification. For the SARS-CoV-2 spike, we synthesized a base construct (HP-GSAS) with residues 1 to 1208 from the Wuhan-Hu-1 strain (GenBank: QHD43416.1) with six stabilizing proline (HexaPro) substitutions at positions 817, 892, 899, 942, 986, and 987 and the S1/S2 furin cleavage site modified to _682_GSAS_685_. Omicron JN.1 and other mutants (JN.1.11, JN.1.11+S31Δ, JN.1.11+Q493E, JN.1.11+S31Δ+Q493E, and KP.3.1.1) were also synthesized using this method. For protein expression, Expi293F cells (Thermo Fisher Scientific: A14527) and FreeStyle 293-F cells (Thermo Fisher Scientific: R79007) were transected with the spike plasmid of interest. Transfected cells were incubated at 37°C with shaking at 110 r.p.m. for 6 days for protein expression. The spike proteins were purified from the supernatants with Strep-Tactin®XT 4Flow® high capacity (IBA Lifescience GmbH: 2-5010-025) and buffer-exchanged to TBS (20 mM Tris and 150 mM NaCl, pH 7.4) before further purification on a Superdex 200 16/90 column (GE Healthcare Biosciences). Protein fractions corresponding to the trimeric spike proteins were collected and concentrated.

### Biolayer interferometry binding assay

BLI assays were performed by using an Octet Red instrument (FortéBio). To measure the binding kinetics of anti-SARS-CoV-2 Fabs and RBDs, the Fabs were diluted with kinetics buffer (1x PBS, pH 7.4, 0.01% BSA and 0.002% Tween 20) to 20 µg/ml. The Fabs were then loaded onto anti-human FAB2G biosensors and interacted with 5-fold gradient dilution (500 nM – 20 nM) of SARS-CoV-2 RBDs. The assay consisted of the following steps. 1) baseline: 1 min with 1x kinetics buffer; 2) loading: 90 seconds with Fabs; 3) wash: 15 seconds wash of unbound Fabs with 1x kinetics buffer; 4) baseline: 1 min with 1x kinetics buffer; 5) association: 90 seconds with RBDs; and 6) dissociation: 90 seconds with 1x kinetics buffer. For estimating K_D_, a 1:1 binding model was used.

To measure the binding kinetics of anti-SARS-CoV-2 Fabs and full-length spikes, twin-strep-tagged spike proteins at 20 μg/mL in 1x kinetics buffer (1x PBS, pH 7.4, 0.01% BSA and 0.002% Tween 20) were loaded onto SA sensors, and then incubated with 12.5, 25, 50, 100, and 200 nM of VYD222 Fab. The assay consisted of five steps: 1) baseline: 60 s with 1x kinetics buffer; 2) loading: 180 s with Twin-strep-tagged spike proteins; 3) baseline: 90 s with 1x kinetics buffer; 4) association: 240 s with VYD222 Fab; and 5) dissociation: 240 s with 1x kinetics buffer. For estimating the K_D_ values, a 1:1 binding model was used.

### Crystallization and structural determination

ADI-77310/RBD-WT, VYD222/RBD-WT and VYD222/RBD-BA.5 complexes were formed by mixing each of the protein components at an equimolar ratio and incubating overnight at 4°C. The protein complex was adjusted to 12 mg/ml and screened for crystallization using the 384 conditions of the JCSG Core Suite (Qiagen) on our robotic CrystalMation system (Rigaku) at Scripps Research. Crystallization trials were set-up by the vapor diffusion method in sitting drops containing 0.1 μl of protein and 0.1 μl of reservoir solution. For the ADI-77310/RBD-WT complex, optimized crystals were grown in drops containing 0.08 M sodium acetate, pH 4.4, 1.45 M ammonium sulfate, and 20% glycerol (v/v) at 20°C. For the VYD222/RBD-WT complex, optimized crystals were grown in drops containing 0.1 M sodium citrate, pH 5.0, and 1.15 M ammonium sulfate at 20°C. Crystals appeared on day 3, were harvested on day 15 by soaking in reservoir solution supplemented with 15% (v/v) ethylene glycol and 20% (v/v) glycerol, respectively, and then flash cooled and stored in liquid nitrogen until data collection. Diffraction data were collected at cryogenic temperature (100 K) at beamline 5.0.1 of the Advanced Light Source (ALS). For the VYD222/RBD-BA.5 complex, crystals were set-up by the vapor diffusion method in sitting drops containing 0.5 μl of protein and 0.5 μl of reservoir solution, and grown in drops containing 0.1 M sodium acetate, pH 4.7, 1.1 M ammonium sulfate, and 0.2 M NaCl at 20°C. Crystals appeared on day 3, were harvested on day 15 by soaking in reservoir solution supplemented with 15% (v/v) ethylene glycol, and then flash cooled and stored in liquid nitrogen until data collection. Diffraction data were collected at cryogenic temperature (100 K) at beamline 23ID-B of the Advanced Photon Source (APS).

Diffraction data were processed with HKL2000 (*33*) for the datasets of ADI-77310/RBD-WT and VYD222/RBD-WT, and with xia2-dials (*34*) for the dataset of VYD222/RBD-BA.5. Structures were solved by molecular replacement using PHASER (*35*). Models for molecular replacement were derived from PDB 7U2D (*1*). Iterative model building and refinement were carried out in COOT (*36*) and PHENIX (*37*), respectively. Epitope and paratope residues, as well as their interactions, were identified by accessing PISA at the European Bioinformatics Institute (http://www.ebi.ac.uk/pdbe/prot_int/pistart.html) (*11*).

### Conservation analysis of SARS-CoV-2 spike proteins

Conservation analysis of SARS-CoV-2 spike proteins was carried out with ConSurf (*38*). The sequence accession codes of SARS-CoV-2 spike proteins are listed as follows: SARS-CoV-2 Wuhan (MN908947), B.1.1.7 (QWE88920), B.1.351 (QRN78347), P.1 (QRX39425), B.1.617.2 (QUD52764), Omicron BA.1 (UFO69279), BA.2 (UJP23605), BA.2.12.1 (UMZ92892), BA.2.75 (UTM82166), BA.4 (UPP14409), BA.4.6 (UOZ45804), BA.5 (UOZ45804), BQ.1.1 (UWM38596), XBB.1.5 (UZG29433), EG.5.1 (WGM84363), JN.1 (WPF38074), KP.3.1.1 (EPI_ISL_19308802), and XEC (EPI_ISL_19283843).

### Pseudovirus neutralization assay

PhenoSense pseudovirus neutralization assays were performed by Monogram Biosciences (Labcorp) as described in Huang et al. 2021 (*39*). Briefly, pseudoviruses bearing SARS-CoV-2 variant spike proteins were produced by co-transfecting HEK293 cells with a codon-optimized spike sequence expression vector and an HIV genomic vector with a firefly luciferase reporter gene replacing the HIV envelope gene. Culture supernatants were harvested 48 hours post transfection, filtered, and frozen at < –70°C. Pseudovirus titers were determined by inoculating HEK293 cells that were transiently transfected with hACE2 and TMPRSS2 expression vectors and measuring luciferase activity in relative light units (RLU) following incubation at 37°C for 3 days. Virus inoculum for the assay was standardized for all variants based on the screening RLUs.

To test antibody neutralization, a predetermined amount of pseudovirus was incubated with titrating amounts of test mAb for 1 hour at 37°C before adding to HEK293 cells expressing hACE2 and TMPRSS2. Pseudovirus infection was allowed to occur for 3 days before cells were assessed for luciferase activity. Luciferase activity was determined by adding Steady Glo (Promega) and measuring luciferase signal (RLU) using a luminometer. Percent neutralization was calculated using the formula 100% x {1 – (RLU virus + sample + cells) / (Avg RLU virus + diluent + cells)}.

### Deep mutational scanning assay

Deep mutational scanning libraries for SARS-CoV-2 RBD variants Wuhan-Hu-1, Omicron BA.2, Omicron XBB.1.5, and Omicron BA.2.86 were described previously, including details of library construction, library availability, and orthogonal measurement of mutational impacts on RBD expression and ACE2-binding affinity (*40–42*). These libraries contain virtually all single amino acid replacements, and in XBB.1.5 and BA.2.86, single-codon-deletions, in each RBD background in a yeast-surface display platform.

Duplicate yeast-display deep mutational scanning libraries were induced for RBD expression, and 5 OD*mL of yeast were incubated for one hour at room temperature in 1 mL solution of mAb at a concentration corresponding to the EC_90_ of the mAb for the respective yeast-displayed wildtype RBD determined from isogenic binding assays. In parallel, controls for FACS gate setting were prepared with 0.5 OD*mL of yeast displaying the wildtype parental control construct incubated in 100 µL of solution of mAb at the matched EC_90_ concentration (1x) or 1/10 this EC_90_ concentration (0.1x). Cells were washed, incubated with 1:100 FITC-conjugated chicken anti-Myc antibody (Immunology Consultants CMYC-45F) and 1:200 PE-conjugated goat anti-human IgG (Jackson ImmunoResearch 109-115-098) to label bound antibody, and washed in preparation for FACS.

Antibody-escape cells in each library were selected via FACS on a BD FACSAria II cell sorter. FACS selection gates were drawn to capture ∼50% of yeast expressing the parental wildtype control labeled at a 0.1x reduced antibody labeling concentration (see representative gating scheme in Figure S4A). For each sample, 5 million RBD+ cells were processed on the sorter while collecting cells in the antibody-escape bin, which were expanded overnight, plasmid purified, and barcodes sequenced on an Illumina NextSeq. In parallel, plasmid samples were purified from ∼30 OD*mL of pre-sorted yeast library cultures and sequenced to establish pre-sort barcode frequences. Barcode reads are available on the NCBI SRA, BioProject PRJNA770094, BioSample SAMN46902828.

Demultiplexed Illumina barcode reads were matched to previously annotated library barcodes and associated with the linked RBD mutant using dms_variants (version 0.8.9), yielding a table of counts of each barcode in each population, available at: https://github.com/tstarrlab/SARS-CoV-2-RBD_Omicron_MAP_VYD222/tree/main/results/counts.

The escape fraction of each barcoded variant was computed from sequencing counts in the pre-sort and antibody-escape populations via the formula:

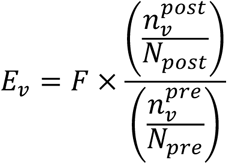

where *F* is the total fraction of the library that escapes antibody binding, *n_v_* is the counts of variant *v* in the pre- or post-sort samples with a pseudocount addition of 0.5, and *N* is the total sequencing count across all variants pre- or post-sort. These escape fractions represent the estimated fraction of cells expressing a particular variant that fall in the escape bin, which scales from 0 for a mutation that never causes sufficient loss of binding to drive cells into the antibody-escape bin, to 1 for a mutation that escapes binding >10-fold such that the variant falls into the antibody-escape bin defined by the 0.1x reduced concentration control labeling gates. We applied computational filters to remove mutants with low pre-selection sequencing counts or highly deleterious mutations that escape antibody binding artefactually due to poor RBD surface expression, specifically mutants with orthogonally measured ACE2-binding impacts of <–3 (1000-fold loss of ACE2 binding) or expression scores of <–1.25, <–0.955, <–1.25, and <–0.75 for Wuhan-Hu-1, BA.2, XBB.1.5, and BA.2.86, respectively, accounting for the variation in baseline expression levels of different wildtype variants and differences in the scaling of arbitrary-unit expression values across deep mutational scanning experiments. Final per-mutant escape fractions were computed as the average escape fraction across barcodes linked to the same mutant within replicates, with the correlation between replicate library selections shown in Figure S4B. Final escape fraction measurements averaged across replicates are available at: https://github.com/tstarrlab/SARS-CoV-2-RBD_Omicron_MAP_VYD222/tree/main/results/supp_data.

## ACKNOWLEDGMENTS

We thank Robyn Stanfield for assistance in data collection. This work was supported by the NIH R01 AI190286 (M.Y., I.A.W.), the Gates Foundation INV-004923 (I.A.W.), Searle Scholars Foundation (T.N.S.), Adagio (grant to I.A.W.), and Invivyd (grant to T.N.S.)

Diffraction data were collected at National ALS and APS synchrotron facilities. The Advanced Light Source is a Department of Energy Office of Science User Facility under Contract No. DE-AC02-05CH11231. The Pilatus detector on 5.0.1. was funded under NIH grant S10OD021832. The ALS-ENABLE beamlines are supported in part by the NIH, National Institute of General Medical Sciences, grant P30 GM124169. This research used resources of the Advanced Photon Source, a U.S. Department of Energy (DOE) Office of Science User Facility operated for the DOE Office of Science by Argonne National Laboratory under contract no. DE-AC02-06CH11357. Extraordinary facility operations were supported, in part, by the DOE Office of Science through the National Virtual Biotechnology Laboratory, a consortium of DOE national laboratories focused on the response to COVID-19, with funding provided by the Coronavirus CARES Act. We thank the Flow Cytometry Core Facility at the University of Utah Health Sciences Campus supported by NIH (5P30CA042014-24), and the University of Utah Center for High Performance Computing supported by NIH (1S10OD021644-01A1).

We would like to thank Emma S. Esterman, Chengzi I. Kaku, Elizabeth McGurk, Elizabeth Parker, Michael E. Brown, C. Garrett Rappazzo, and James C. Geoghegan of Adimab, LLC, for their contribution to the efforts leading to the derivation of VYD222.

## AUTHOR CONTRIBUTIONS

Conceptualization: M.Y., B.R.W., L.W., R.A., I.A.W.

Methodology: M.Y., B.R.W., L.W., T.N.S., RA.

Investigation: M.Y., Z.F., W.B.F., A.L.T., T.N.S., B.R.W., C.P., X.Y., G.W.

Visualization: M.Y., Z.F., B.R.W., T.N.S.

Funding acquisition: M.Y., T.N.S., I.A.W.

Project administration: B.R.W., T.N.S., I.A.W.

Supervision: B.R.W., T.N.S., I.A.W.

Writing – original draft: M.Y., B.R.W., I.A.W.

Writing – review & editing: All authors

## COMPETING INTERESTS

Some of the authors were employees of Invivyd, Inc. (B.W., C.P., L.W., and R.A.) at the time this research was conducted and may hold stock or shares in Invivyd, Inc. L.W. is currently an employee of Moderna, Inc. Other authors declare that they have no competing interests.

## DATA AVAILABILITY

The X-ray coordinates and structure factors have been deposited to the RCSB Protein Data Bank under accession codes: 9PMU, 9PMV, and 9PMX. DMS sequencing data are available from the NCBI SRA, BioProject PRJNA770094, BioSample SAMN46902828.

