## Supplemental figures tables for "Structural and functional analysis of VYD222: a broadly neutralizing antibody against SARS-CoV-2 variants"

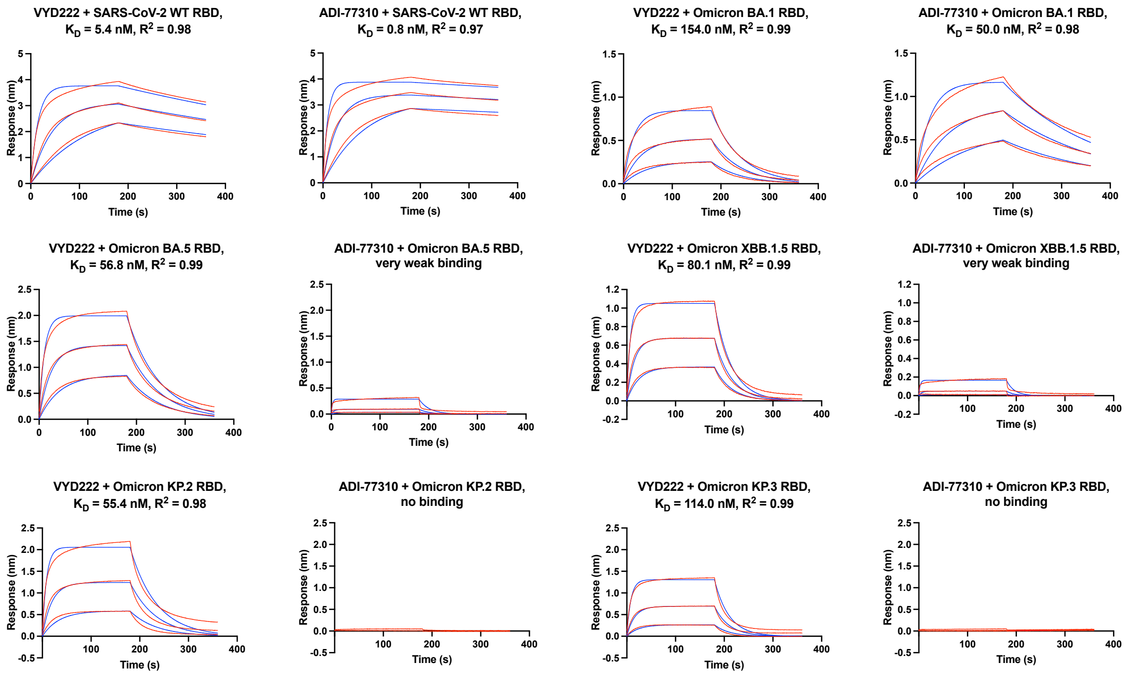


**Figure S1. Sensorgrams for binding of Fabs to RBDs of SARS-CoV-2 measured by biolayer interferometry (BLI).** X-axis represents time of reaction (s). Y-axis represents the response. Blue lines represent the response curves and red lines represent the 1:1 binding model. Binding kinetics were measured for the RBDs at 5-fold dilution ranging from 500 nM to 20 nM.

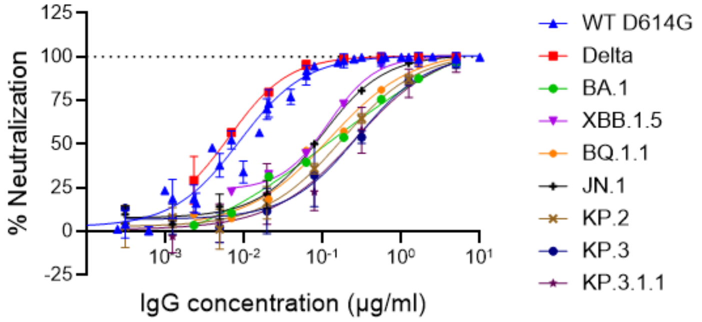


**Figure S2. Dose-response curves in pseudovirus neutralization assay for VYD222.** Percent neutralization values obtained from the PhenoSense assay for VYD222 against the indicated variant were curve fitted using a 4PL nonlinear regression model (GraphPad Prism version 10.1.2).

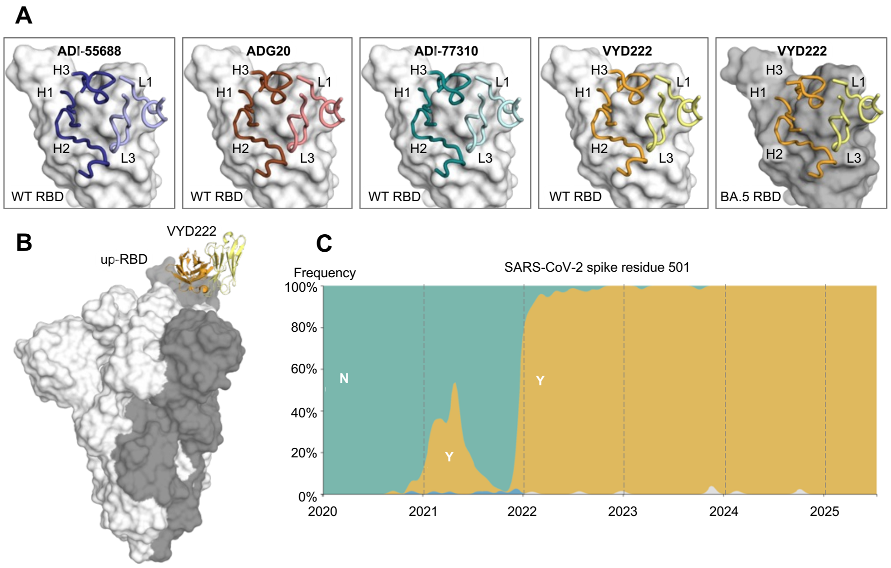


**Figure S3. Interactions between SARS-CoV-2 spike and ADI-55688, ADG20, ADI-77310, and VYD222. (A)** WT and BA.5 RBDs are represented by white and dark grey surfaces. CDRs that interact with SARS-CoV-2 are shown as tubes. **(B)** VYD222/RBD complex structure superimposed onto a full spike protein (PDB 8G71). The protomers with RBDs in down conformations are in white, while that with an RBD in the up position is in dark grey. **(C)** Frequency of amino acids at position 501 on SARS-CoV-2 spike as of July 2025. Data were sourced from nextstrain.org (GISAID data) (*1, 2*).

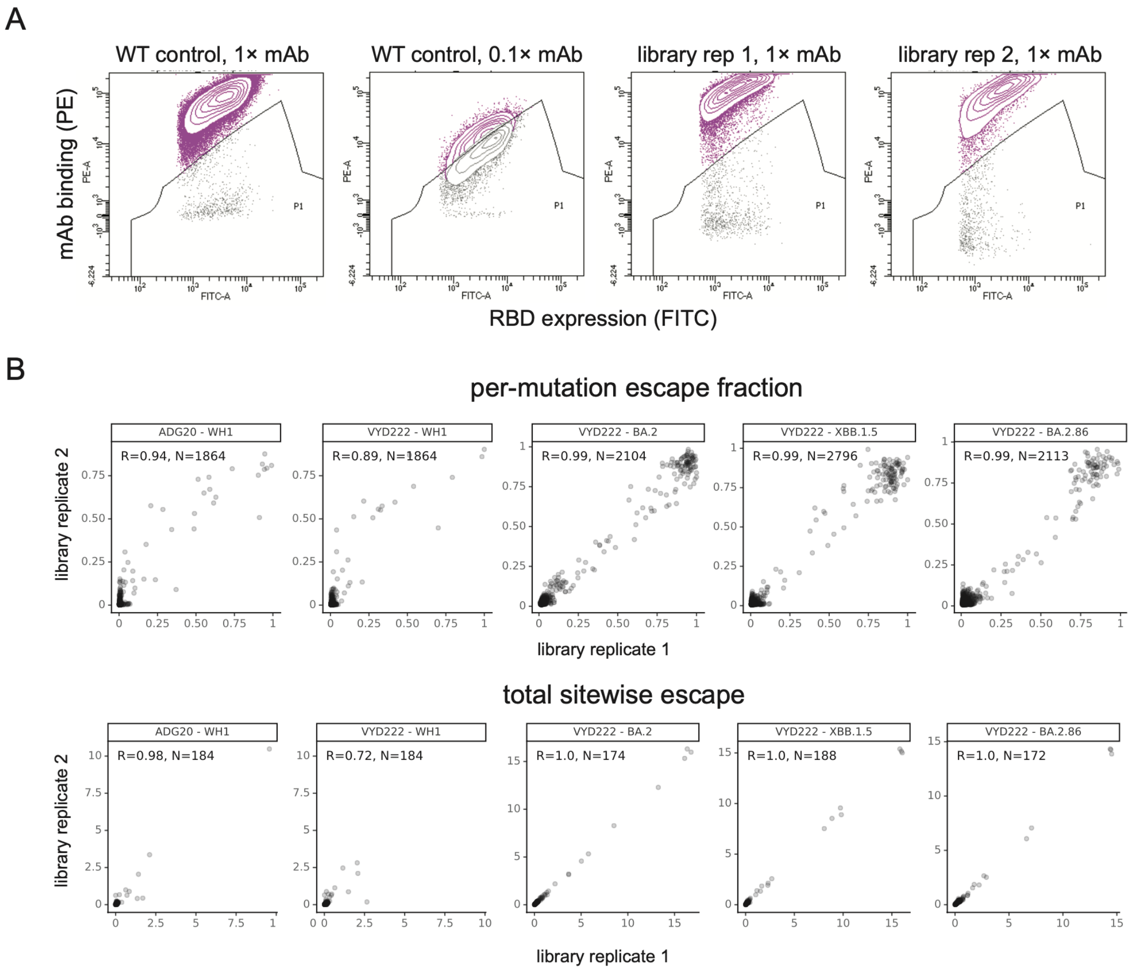


**Figure S4. ADG20 or VYD222 binding to SARS-CoV-2 variant deep mutational scanning libraries. (A)** Representative FACS gates used to identify mutations that escape antibody binding. An antibody-escape gate is drawn that captures approximately 50% of the cells in the respective wildtype control labeled at 0.1x the library selection antibody concentration. The "escape fraction" determined from deep sequencing of pre- and post-sort populations estimates the fraction of yeast cells expressing a particular mutant genotype that fell into this antibody-escape FACS gate. **(B)** For each experiment, the correlation in the per-mutation escape fraction (top) or total escape per site (sum of per-mutation escape fractions; bottom) between duplicate library selections.

| | 403 | 404 | 405 | 406 | 408 | 409 | 436 | 437 | 439 | 493 | 500 | 501 | 502 | 503 | 504 | 505 | 506 |
|---|---|---|---|---|---|---|---|---|---|---|---|---|---|---|---|---|---|
| SARS-CoV-2_MN908947 | R | G | D | V | R | Q | W | N | N | Q | T | N | G | V | G | Y | Q |
| alpha | R | G | D | V | R | Q | W | N | N | Q | T | Y | G | V | G | Y | Q |
| beta | R | G | D | V | R | Q | W | N | N | Q | T | Y | G | V | G | Y | Q |
| gamma | R | G | D | V | R | Q | W | N | N | Q | T | Y | G | V | G | Y | Q |
| mu | R | G | D | V | R | Q | W | N | N | Q | T | Y | G | V | G | Y | Q |
| delta | R | G | D | V | R | Q | W | N | N | Q | T | N | G | V | G | Y | Q |
| KP.3.1.1 | K | G | N | V | S | Q | W | N | N | E | T | Y | G | V | G | H | Q |
| KP.1.1 | K | G | N | V | S | Q | W | N | N | Q | T | Y | G | V | G | H | Q |
| KP.2 | K | G | N | V | S | Q | W | N | N | Q | T | Y | G | V | G | H | Q |
| JN.1 | K | G | N | V | S | Q | W | N | N | Q | T | Y | G | V | G | H | Q |
| BA.2.86 | K | G | N | V | S | Q | W | N | N | Q | T | Y | G | V | G | H | Q |
| EG.5.1 | R | G | N | V | S | Q | W | N | N | Q | T | Y | G | V | G | H | Q |
| XBB.1.5 | R | G | N | V | S | Q | W | N | N | Q | T | Y | G | V | G | H | Q |
| XBB.1.16 | R | G | N | V | S | Q | W | N | N | Q | T | Y | G | V | G | H | Q |
| BA.1 | R | G | D | V | R | Q | W | N | N | R | T | Y | G | V | G | H | Q |
| BQ.1.1 | R | G | N | V | S | Q | W | N | N | Q | T | Y | G | V | G | H | Q |
| BA.2 | R | G | N | V | S | Q | W | N | N | R | T | Y | G | V | G | H | Q |
| BA.4/5 | R | G | N | V | S | Q | W | N | N | Q | T | Y | G | V | G | H | Q |
| Rs4084_KY417144 | K | G | D | V | R | Q | W | N | N | R | T | A | G | V | G | H | Q |
| Rs4231_KY417146 | K | G | D | V | R | Q | W | N | N | R | T | A | G | V | G | H | Q |
| RsSHC014_KC881005 | K | G | D | V | R | Q | W | N | N | R | T | A | G | V | G | H | Q |
| LYRa11_KF569996 | K | G | D | V | R | Q | W | N | R | N | T | N | G | I | G | Y | Q |
| WIV16_KT444582 | K | G | D | V | R | Q | W | N | R | N | T | N | G | I | G | Y | Q |
| WIV1_KF367457 | K | G | D | V | R | Q | W | N | R | N | T | N | G | I | G | Y | Q |
| Rs7327_KY417151 | K | G | D | V | R | Q | W | N | R | N | T | N | G | I | G | Y | Q |
| SARSCoV1_Sin852_HP03L_AY559082 | K | G | D | V | R | Q | W | N | R | N | T | T | G | I | G | Y | Q |
| SARSCoV1_PC413_PC04_AY613948 | K | G | D | V | R | Q | W | N | R | R | T | T | G | I | G | Y | Q |
| SARSCoV1_SZ3_PC03_AY304486 | K | G | D | V | R | Q | W | N | R | K | T | T | G | I | G | Y | Q |
| Pangolin_GXP2V_EPI_ISL_410542 | K | G | D | V | R | Q | W | N | V | E | T | T | G | V | N | Y | Q |
| RaTG13_MN996532 | T | G | D | V | R | Q | W | N | K | Y | T | D | G | V | G | H | Q |
| Pangolin_GDconsensus_Lam2020 | R | G | D | V | R | Q | W | N | N | Q | T | N | G | V | G | Y | Q |
| BtKY72_KY352407 | K | G | D | V | R | Q | W | N | N | K | T | V | G | V | G | Y | Q |

**Figure S5. Sequence alignment of VYD222 epitope residues in SARS-CoV-2 variants and other sarbecoviruses.** ACE2-binding sarbecoviruses were identified in Starr et al. 2022 (*3*) and updated with recently emerged SARS-CoV-2 variants.

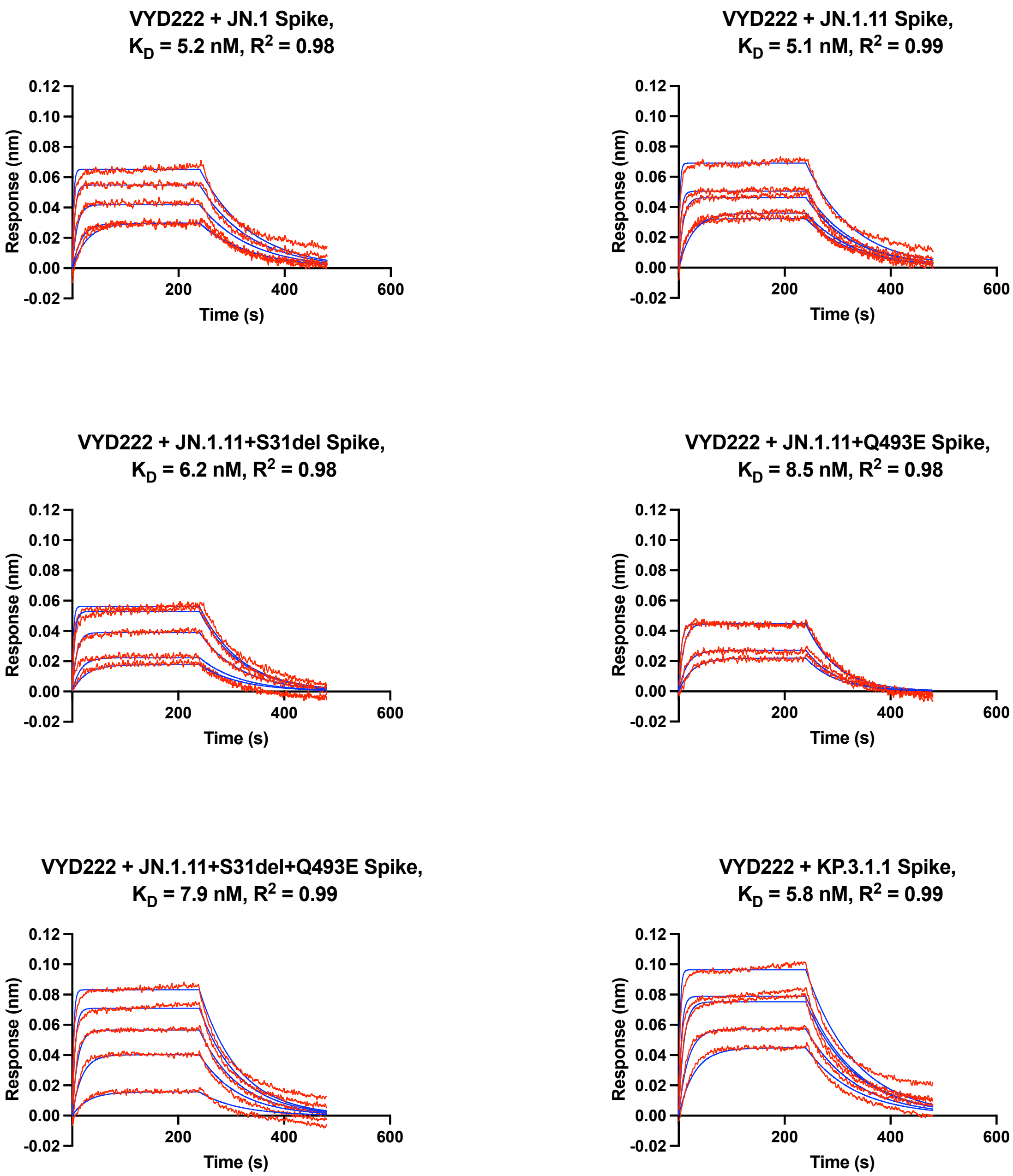


**Figure S6. Sensorgrams for binding of Fabs to SARS-CoV-2 spikes measured by BLI.** X-axis represents time of reaction (s). Y-axis represents the response. Blue lines represent the response curves and purple lines represent the 1:1 binding model. Binding kinetics were measured for the Fabs at 2-fold dilution ranging from 200 nM to 12.5 nM.

### Table S1. X-ray data collection and refinement statistics

| **Data collection** | ADI-77310 + SARS-CoV-2 RBD WT | VYD222 + SARS-CoV-2 RBD WT | VYD222 + SARS-CoV-2 RBD BA.5 |
|---|---|---|---|
| Beamline | ALS5.0.1 | ALS5.0.1 | APS23ID-B |
| Wavelength (Å) | 0.9774 | 0.9774 | 1.033 |
| Space group | $P\ 4_1$ | $P\ 4_1$ | $P\ 4_1$ |
| Unit cell parameters | | | |
| a, b, c (Å) | 100.3, 100.3, 79.8 | 100.0, 100.0, 78.9 | 142.4, 142.4, 81.4 |
| α, β, γ (°) | 90, 90, 90 | 90, 90, 90 | 90, 90, 90 |
| Resolution (Å) [a] | 50.0-3.06 | 50.0-3.00 | 71.2-2.70 |
| Unique reflections [a] | 15,127 (722) | 15,708 (774) | 44,976 (2,237) |
| Redundancy [a] | 13.4 (11.8) | 13.2 (11.2) | 11.8 (12.0) |
| Completeness (%) [a] | 100 (99.9) | 99.9 (99.1) | 100 (99.1) |
| $\langle I/\sigma_I \rangle$ [a] | 9.3 (0.8) | 11.0 (0.9) | 11.4 (0.7) |
| $R_{sym}$[b] (%) [a] | 27.5 (>100) | 19.5 (>100) | 15.9 (>100) |
| $R_{pim}$[b] (%) [a] | 7.8 (64.8) | 5.5 (46.6) | 4.8 (48.2) |
| $CC_{1/2}$[c] (%) [a] | 99.0 (45.8) | 99.7 (61.4) | 99.4 (77.6) |
| **Refinement statistics** | | | |
| Resolution (Å) | 44.9-3.06 | 50.0-3.00 | 63.7-2.70 |
| Reflections (work) | 14,677 | 15,386 | 44,910 |
| Reflections (test) | 708 | 1,311 | 2,006 |
| $R_{cryst}$[d] / $R_{free}$[e] (%) | 23.4/26.3 | 22.1/26.3 | 20.1/25.0 |
| No. of copies per ASU | 1 | 1 | 2 |
| No. of atoms | 4,852 | 4,813 | 9,722 |
| RBD | 1,560 | 1,550 | 3,129 |
| Fab | 3,246 | 3,234 | 6,476 |
| Ligands [f] | 30 | 15 | 51 |
| Solvent | 16 | 14 | 66 |
| Average *B*-values ($Å^2$) | 59 | 67 | 77 |
| RBD | 58 | 65 | 96 |
| Fab | 60 | 68 | 67 |
| Ligands | 77 | 89 | 106 |
| Solvent | 48 | 41 | 59 |
| Wilson *B*-value ($Å^2$) | 72 | 71 | 67 |
| **RMSD from ideal geometry** | | | |
| Bond length (Å) | 0.004 | 0.003 | 0.005 |
| Bond angle (°) | 0.56 | 0.53 | 0.70 |
| **Ramachandran statistics (%)** [g] | | | |
| Favored | 95.5 | 95.5 | 94.9 |
| Outliers | 0.16 | 0.16 | 0.16 |
| **PDB code** | 9PMU | 9PMV | 9PMX |

[a] Numbers in parentheses refer to the highest resolution shell.
[b] $R_{sym} = \Sigma_{hkl} \Sigma_i | I_{hkl,i} - \langle I_{hkl} \rangle | / \Sigma_{hkl} \Sigma_i I_{hkl,i}$ and $R_{pim} = \Sigma_{hkl} (1/(n-1))^{1/2} \Sigma_i | I_{hkl,i} - \langle I_{hkl} \rangle | / \Sigma_{hkl} \Sigma_i I_{hkl,i}$, where $I_{hkl,i}$ is the scaled intensity of the $i^{th}$ measurement of reflection h, k, l, $\langle I_{hkl} \rangle$ is the average intensity for that reflection, and *n* is the redundancy.
[c] $CC_{1/2}$ = Pearson correlation coefficient between two random half datasets.
[d] $R_{cryst} = \Sigma_{hkl} | F_o - F_c | / \Sigma_{hkl} | F_o | \times 100$, where $F_o$ and $F_c$ are the observed and calculated structure factors, respectively.
[e] $R_{free}$ was calculated as for $R_{cryst}$, but on a test set comprising 5%–8.5% of the data excluded from refinement.
[f] Bound ligands are $SO_4$ and acetate.
[g] From MolProbity (*4*).

### Table S2. BSA values of published antibodies targeting SARS-CoV-2 RBD as calculated by PISA (5)

| Name | PDB | Epitope | BSA (Å²) |
|---|---|---|---|
| CR3022 | 6W41 | RBD | 1024 |
| CC12.1 | 6XC2 | RBD | 1347 |
| CC12.3 | 6XC4 | RBD | 885 |
| CV30 | 6XE1 | RBD | 1036 |
| Fab2-4 | 6XEY | RBD | 754 |
| CV07-270 | 6XKP | RBD | 817 |
| CV07-250 | 6XKQ | RBD | 921 |
| HbnC3t1p1_C6 | 7B0B | RBD | 616 |
| COVOX-316 | 7BEH | RBD | 740 |
| COVOX-150 | 7BEI | RBD | 1241 |
| COVOX-158 | 7BEJ | RBD | 1131 |
| COVOX-269 | 7BEM | RBD | 1349 |
| COVOX-253 | 7BEN | RBD | 749 |
| COVOX-384 | 7BEP | RBD | 738 |
| ION-300 | 7BNV | RBD | 676 |
| P2B-2F6 | 7BWJ | RBD | 649 |
| B38 | 7BZ5 | RBD | 1222 |
| CB6 | 7C01 | RBD | 1087 |
| P2C-1F11 | 7CDI | RBD | 958 |
| P2C-1A3 | 7CDJ | RBD | 922 |
| BD-604 | 7CH4 | RBD | 1150 |
| BD-629 | 7CH5 | RBD | 1057 |
| BD-236 | 7CHB | RBD | 1135 |
| BD-368-2 | 7CHC | RBD | 970 |
| P5A-1D2 | 7CHO | RBD | 1013 |
| P5A-3C8 | 7CHP | RBD | 1273 |
| P22A-1D1 | 7CHS | RBD | 1141 |
| P4A1 | 7CJF | RBD | 1189 |
| P2B-1A1 | 7CZP | RBD | 1117 |
| P2B-1A10 | 7CZQ | RBD | 1351 |
| P5A-1B8 | 7CZR | RBD | 1006 |
| P5A-2G9 | 7CZT | RBD | 1234 |
| P5A-1B6 | 7CZU | RBD | 960 |
| P5A-2G7 | 7CZW | RBD | 745 |
| P5A-1B9 | 7CZX | RBD | 890 |
| P5A-2F11 | 7CZY | RBD | 722 |
| P5A-3A1 | 7D0C | RBD | 860 |
| Ab_58G6 | 7E3L | RBD | 689 |
| BD-623 | 7E7Y | RBD | 634 |
| BD-503 | 7EK0 | RBD | 1178 |
| GW01 | 7EPX | RBD | 825 |
| BD-813 | 7EY0 | RBD | 920 |
| BD-667 | 7EY4 | RBD | 1204 |
| BD-821 | 7EY5 | RBD | 592 |
| BD-804 | 7EYA | RBD | 1085 |
| JS026 | 7F7E | RBD | 716 |
| PD36-5D2 | 7FAE | RBD | 798 |
| T6 | 7FJO | RBD | 718 |
| COVA2-04 | 7JMO | RBD | 1191 |
| COVA2-39 | 7JMP | RBD | 672 |
| COVA1-16 | 7JMW | RBD | 827 |
| S2H13 | 7JV2 | RBD | 701 |
| S2A4 | 7JVA | RBD | 856 |
| S2H14 | 7JX3 | RBD | 1070 |
| S2-M11 | 7K43 | RBD | 657 |
| C102 | 7K8M | RBD | 1069 |
| C002 | 7K8S | RBD | 801 |
| C119 | 7K8W | RBD | 801 |
| C144 | 7K90 | RBD | 800 |
| C1A-B3 | 7KFV | RBD | 1206 |
| C1A-B12 | 7KFW | RBD | 1123 |
| C1A-C2 | 7KFX | RBD | 1171 |
| C1A-F10 | 7KFY | RBD | 1141 |
| Bamlanivimab | 7KMG | RBD | 879 |
| LY-CoV488 | 7KMH | RBD | 903 |
| LY-CoV481 | 7KMI | RBD | 1126 |
| Fab2-43 | 7L56 | RBD | 755 |
| Fab2-15 | 7L5B | RBD | 893 |
| AZD-8895 | 7L7D | RBD | 647 |
| AZD-1061 | 7L7E | RBD | 757 |
| DH1041 | 7LAA | RBD | 721 |
| DH1047 | 7LD1 | RBD | 806 |
| DH1043 | 7LJR | RBD | 1089 |
| CV38-142 | 7LM8 | RBD | 862 |
| CV05-163 | 7LOP | RBD | 1045 |
| CV503 | 7LQ7 | RBD | 1054 |
| Fab1-57 | 7LS9 | RBD | 784 |
| Fab2-7 | 7LSS | RBD | 768 |
| CV2-75 | 7M3I | RBD | 809 |
| BG4-25 | 7M6D | RBD | 1120 |
| BG10-19 | 7M6E | RBD | 1018 |
| BG7-15 | 7M6G | RBD | 755 |
| Zhou_47D1 | 7MF1 | RBD | 707 |
| B1-182-1 | 7MLZ | RBD | 795 |
| LY-CoV1404 | 7MMO | RBD | 759 |
| PDI-37 | 7MZF | RBD | 1163 |
| PDI-42 | 7MZG | RBD | 1131 |
| PDI-93 | 7MZJ | RBD | 1212 |
| PDI-96 | 7MZK | RBD | 571 |
| PDI-210 | 7MZL | RBD | 1128 |
| PDI-215 | 7MZM | RBD | 1046 |
| PDI-231 | 7MZN | RBD | 1178 |
| WRAIR-2057 | 7N4I | RBD | 839 |
| WRAIR-2173 | 7N4J | RBD | 937 |
| WRAIR-2125 | 7N4L | RBD | 673 |
| Fab2-36 | 7N5H | RBD | 914 |
| ION-360 | 7NP1 | RBD | 876 |
| COVOX-222 | 7NX6 | RBD | 1025 |
| COVOX-278 | 7OR9 | RBD | 841 |
| P5C3 | 7P40 | RBD | 414 |
| FD-11A | 7PQZ | RBD | 781 |
| FD-5D | 7PR0 | RBD | 844 |
| Beta-6 | 7PRY | RBD | 626 |
| Beta-22 | 7PRZ | RBD | 913 |
| Beta-24 | 7PS0 | RBD | 761 |
| Beta-27 | 7PS1 | RBD | 988 |

| Name | PDB | Epitope | BSA (Å²) |
|---|---|---|---|
| Beta-38 | 7PS4 | RBD | 688 |
| Beta-47 | 7PS5 | RBD | 826 |
| FI-3A | 7Q0G | RBD | 957 |
| Beta-49 | 7Q6E | RBD | 944 |
| Beta-50 | 7Q9F | RBD | 954 |
| CV2.1169 | 7QEZ | RBD | 690 |
| CV2.3235 | 7QF0 | RBD | 1349 |
| CV2.6264 | 7QF1 | RBD | 761 |
| Beta-55 | 7QNW | RBD | 1037 |
| COVOX-58 | 7QNY | RBD | 1344 |
| P2G3 | 7QTI | RBD | 604 |
| C051 | 7R8N | RBD | 941 |
| C548 | 7R8O | RBD | 928 |
| C022 | 7RKU | RBD | 973 |
| C118 | 7RKV | RBD | 1022 |
| PDI-222 | 7RR0 | RBD | 861 |
| CC6.33 | 7RU3 | RBD | 847 |
| CC6.30 | 7RU5 | RBD | 1065 |
| CS23 | 7S5P | RBD | 1084 |
| CS44 | 7S5Q | RBD | 901 |
| CV07-287 | 7S5R | RBD | 631 |
| J08 | 7S6J | RBD | 735 |
| R40-1G8 | 7SC1 | RBD | 1115 |
| Liu_10-40 | 7SD5 | RBD | 979 |
| Liu_10-28 | 7SI2 | RBD | 949 |
| 54042-4 | 7T01 | RBD | 592 |
| GAR5 | 7T72 | RBD | 1083 |
| ADI-62113 | 7T7B | RBD | 874 |
| S2K146 | 7TAS | RBD | 985 |
| A19-61-1 | 7TB8 | RBD | 477 |
| DH1042 | 7THE | RBD | 983 |
| 002-02 | 7U0Q | RBD | 1084 |
| 002-13 | 7U0X | RBD | 853 |
| ADI-55688 | 7U2E | RBD | 657 |
| NE12 | 7U9O | RBD | 874 |
| NA8 | 7U9P | RBD | 1003 |
| CoV11 | 7URQ | RBD | 1161 |
| XG014 | 7V2A | RBD | 869 |
| TAU-2303 | 7WBZ | RBD | 1204 |
| CZ-D7 | 7WCH | RBD | 727 |
| XGv282 | 7WE7 | RBD | 735 |
| XGv289 | 7WE9 | RBD | 581 |
| XGv347 | 7WEA | RBD | 741 |
| XGv265 | 7WED | RBD | 753 |
| ZB8 | 7WH8 | RBD | 1049 |
| 6M6 | 7WJY | RBD | 1208 |
| NCV2SG53 | 7WN2 | RBD | 906 |
| NCV2SG48 | 7WNB | RBD | 1265 |
| 553-15 | 7WO4 | RBD | 864 |
| 553-60 | 7WOA | RBD | 716 |
| 553-49 | 7WOG | RBD | 1138 |
| BD55-3152 | 7WR8 | RBD | 895 |
| BD55-4637 | 7WRJ | RBD | 1084 |
| Ab_510A5 | 7WS4 | RBD | 748 |
| BD55-3500 | 7WSC | RBD | 923 |
| XGv051 | 7WTF | RBD | 907 |
| XGv264 | 7WTH | RBD | 824 |
| XGv-264 | 7WTI | RBD | 632 |
| XGv286 | 7WTJ | RBD | 558 |
| Ab_55A8 | 7WWI | RBD | 727 |
| ZWD12 | 7WWL | RBD | 696 |
| ZWC6 | 7WWM | RBD | 747 |
| 2G1 | 7X08 | RBD | 631 |
| UT28K | 7X7O | RBD | 648 |
| Ab354 | 7X8W | RBD | 743 |
| Ab159 | 7X8Y | RBD | 579 |
| Ab326 | 7X90 | RBD | 1011 |
| Ab496 | 7X91 | RBD | 981 |
| Ab445 | 7X92 | RBD | 1042 |
| ZCB11 | 7XH8 | RBD | 804 |
| STS165 | 7XIC | RBD | 1090 |
| P2S-2E9 | 7XSA | RBD | 744 |
| P5S-3B11 | 7XSB | RBD | 749 |
| P5S-2B10 | 7XSC | RBD | 908 |
| BD55-1403 | 7Y0C | RBD | 973 |
| BD55-5549 | 7Y0V | RBD | 827 |
| BD55-5514 | 7Y0W | RBD | 899 |
| FP-12A | 7YCK | RBD | 881 |
| IS-9A | 7YCL | RBD | 863 |
| IY-2A | 7YCN | RBD | 1138 |
| P3E6 | 7YKJ | RBD | 708 |
| XG2v024 | 7YR1 | RBD | 872 |
| THSC20.HVTR26 | 7Z0X | RBD | 634 |
| THSC20.HVTR04 | 7Z0Y | RBD | 665 |
| Omi-3 | 7ZF3 | RBD | 1008 |
| Omi-25 | 7ZFD | RBD | 806 |
| Omi-42 | 7ZR7 | RBD | 666 |
| Omi-38 | 7ZR8 | RBD | 945 |
| Pushparaj_Fab47 | 8A94 | RBD | 984 |
| BA.2-10 | 8BBN | RBD | 994 |
| BA.2-36 | 8BBO | RBD | 936 |
| BA.2-23 | 8BCZ | RBD | 1113 |
| S728-1157 | 8D0Z | RBD | 1157 |
| AZ090 | 8DAD | RBD | 857 |
| P1D9 | 8DWA | RBD | 964 |
| P2B4 | 8DXS | RBD | 688 |
| GAR12 | 8DXT | RBD | 1035 |
| GAR3 | 8DXU | RBD | 1105 |
| S2X324 | 8ERQ | RBD | 785 |
| Hastie_1C3 | 8F0G | RBD | 602 |
| YB9-258 | 8HC2 | RBD | 1186 |
| YB13-292 | 8HC8 | RBD | 837 |
| **ADI-77310** | **this study** | **RBD** | **718** |
| **VYD222** | **this study** | **RBD** | **701** |
| **Average** | | | **904** |
| **Standard deviation** | | | **151** |
